# Evaluating the role of reference-genome phylogenetic distance on evolutionary inference

**DOI:** 10.1101/2021.03.03.433733

**Authors:** Aparna Prasad, Eline D Lorenzen, Michael V Westbury

## Abstract

When a high-quality genome assembly of a target species is unavailable, an option to avoid the costly *de novo* assembly process is a mapping-based assembly. However, mapping shotgun data to a distant relative may lead to biased or erroneous evolutionary inference. Here, we used short-read data from a mammal and a bird species (beluga and rowi kiwi) to evaluate whether reference genome phylogenetic distance can impact downstream demographic (PSMC) and genetic diversity (heterozygosity, runs of homozygosity) analyses. We mapped to assemblies of species of varying phylogenetic distance (conspecific to genome-wide divergence of >7%), and *de novo* assemblies created using cross-species scaffolding. We show that while reference genome phylogenetic distance has an impact on demographic analyses, it is not pronounced until using a reference genome with >3% divergence from the target species. When mapping to cross-species scaffolded assemblies, we are unable to replicate the original beluga demographic analyses, but can with the rowi kiwi, presumably reflecting the more fragmented nature of the beluga assemblies. As for genetic diversity estimates, we find that increased phylogenetic distance has a pronounced impact; heterozygosity estimates deviate incrementally as phylogenetic distance increases. Moreover, runs of homozygosity are removed when mapping to any non-conspecific assembly. However, these biases can be reduced when mapping to a cross-species scaffolded assembly. Taken together, our results show that caution should be exercised when selecting the reference genome for mapping assemblies. Cross-species scaffolding may offer a way to avoid a costly, traditional *de novo* assembly, while still producing robust, evolutionary inference.

## Introduction

The large extent of genetic information within the nuclear genome enables powerful evolutionary inferences using just a single individual. Two options are available for genome assembly: mapping-based assemblies using a closely-related species as reference, or *de novo* assemblies. In the former approach, relatively little time and monetary expense is invested in sequencing one individual to high coverage (>20x). After assembly, it is possible to make population-wide evolutionary inferences of the target species, including levels of genetic diversity and inbreeding, adaptive genomic changes, and demographic history (Barnett et al., 2020; Lord et al., 2020; Michael V. Westbury, Petersen, Garde, Heide-Jørgensen, & Lorenzen, 2019).

Although mapping-based assemblies are less costly than *de novo* assemblies, there are some caveats. Biases towards the reference genome allele may be an issue when analysing population-level datasets. Such errors can arise during variant calling, when the alternative allele fails to be called altogether, or when heterozygous sites are incorrectly called as homozygous for the reference allele (Brandt et al., 2015; Ros-Freixedes et al., 2018). Although such issues are known to occur when mapping to a conspecific from a different population, biases caused by mapping to phylogenetically more distant taxa have only somewhat been addressed (Armstrong et al., 2020; M. V. Westbury et al., 2021). Problems with correctly identifying variants may arise due to decreased mapping efficiency as reference-genome phylogenetic distance increases (Shapiro & Hofreiter, 2014). However, the consequences of this on downstream analyses have yet to be comprehensively assessed. This leads to the question of whether the potentially costly *de novo* assembly process can be avoided when assemblies from phylogenetically more distant species are available. Insights into this will be especially important for the study of extinct species, where a conspecific reference genome is unlikely to be available (Barnett et al., 2020; Palkopoulou et al., 2018).

Here, we investigate the influence of the reference genome’s phylogenetic distance to the target species on downstream evolutionary analyses. Specifically, we focused on estimates of (i) demographic history (reconstructed using PSMC), (ii) genetic diversity (genome-wide heterozygosity estimated using ANGSD, ROHan, SAMtools/BCFtools), (iii) inbreeding/runs of homozygosity (using ROHan). Additionally, we investigated whether biases can be overcome by using cross-species scaffolded con-specific assemblies as mapping reference. We applied our methodology to two taxonomically disparate datasets; one based on mammals (beluga and incrementally divergent cetacean species), and one based on birds (rowi kiwi and incrementally divergent paleognath species). We selected these datasets based on the assumption that mammal and bird genomes may respond differently to reference biases, allowing for more generalised conclusions. Furthermore, while beluga whales are a relatively abundant species (Hobbs et al., 2019), rowi kiwi are threatened with extinction and have much lower population numbers (Robertson et al., 2017), which may play a role.

## Materials and methods

A simplified version of the methodologies implemented in this manuscript are presented in figure 1.

**Figure 1:**
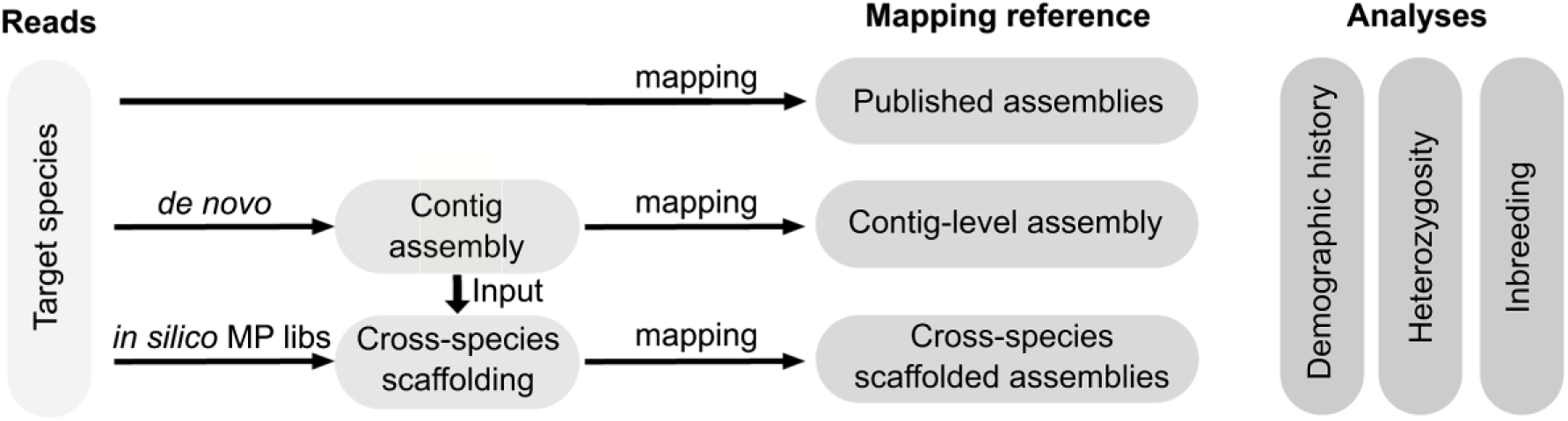
Overview of the approaches used to investigate the role reference genome plays in downstream demographic history and genetic diversity results. Raw reads are mapped to published assemblies, a *de novo* contig-level assembly, or cross-species scaffolded assemblies. Contig-level assemblies are constructed using the raw reads. Cross-species scaffolded assemblies are made by scaffolding the contig assembly using *in-silico* mate-pair (MP) libraries.

### Data

For the cetacean comparative dataset, we downloaded the raw Illumina reads and an assembled genome of the beluga (*Delphinapterus leucas*, Genbank accession code: GCF_002288925.2, SRA code: SRR5197961). In addition, we downloaded genome assemblies for five other cetacean species with varying genomic distance to the beluga (Table 1): narwhal (*Monodon monoceros*, Genbank accession code: GCF_005190385.1), narrow-ridged finless porpoise (*Neophocaena asiaeorientalis*, Genbank accession code: GCF_003031525.1), bottlenose dolphin (*Tursiops truncatus*, Genbank accession code: GCF_001922835.1), sperm whale (*Physeter macrocephalus*, Genbank accession code: GCA_002837175.2), and minke whale (*Balaenoptera acutorostrata*, Genbank accession code: GCF_000493695.1). Assembly length, N50, and level of missing data for each assembly are listed in supplementary table S1.

**Table 1:** Genome-wide pairwise divergence estimates of the species used in this study.

| Species | Compared to | Divergence |
| --- | --- | --- |
| Narwhal | Beluga | 0.0050 |
| Finless porpoise | Beluga | 0.0143 |
| Bottlenose dolphin | Beluga | 0.0202 |
| Sperm whale | Beluga | 0.0318 |
| Minke whale | Beluga | 0.0344 |
| Brown kiwi | Rowi kiwi | 0.0031 |
| Spotted kiwi | Rowi kiwi | 0.0079 |
| Emu | Rowi kiwi | 0.0734 |

For the paleognath comparative dataset, we downloaded raw reads and an assembled genome of the rowi kiwi (Genbank accession: GCF_003343035.1, SRA accession: SRR6918118). We also downloaded published assemblies for three palaeognathae species of varying genomic distance to the rowi kiwi (Table 1): North Island brown kiwi (termed brown kiwi here, *A. mantelli*, Genbank accession: GCF_001039765.1), great spotted kiwi (termed spotted kiwi here, *A. haastii*, Genbank accession: GCA_003342985.1), and emu (*Dromaius novaehollandiae*, Genbank accession: GCF_003342905.1). Assembly length, N50, and levels of missing data for each assembly are listed in supplementary table S2.

### *De novo* and mapping assemblies

All mappings were performed following the same procedure for the beluga/cetacean and rowi kiwi/palaeognathae species datasets. We trimmed adapter sequences and removed reads shorter than 30bp from the raw reads using skewer v0.2.2 (Jiang, Lei, Ding, & Zhu, 2014)), and mapped the trimmed reads using BWA v0.7.15 (Li & Durbin, 2009) and the mem algorithm utilising default parameters. We parsed the output and removed duplicates and reads with a mapping quality lower than 30 with SAMtools v1.6 (Li et al., 2009).

For the beluga, we mapped the beluga short-read data to (i) published assemblies (beluga, narwhal, finless porpoise, bottlenose dolphin, sperm whale, minke whale), (ii) a *de novo* contig-level beluga assembly constructed for the purposes of this study, and (iii) five *de novo* beluga assemblies produced by cross-species scaffolding the contig-level assembly with each published non-beluga assembly using in-silico mate pair (MP) libraries.

For the rowi kiwi, we mapped the rowi short-read data to (i) published assemblies (rowi kiwi, brown kiwi, spotted kiwi, emu), (ii) a *de novo* contig-level rowi kiwi assembly constructed for the purposes of this study, and (iii) three *de novo* rowi kiwi assemblies produced by cross-species scaffolding the contig-level assembly with each of the non-rowi kiwi published assembles using in-silico MP libraries.

We constructed *de novo* contig-level assemblies for both the beluga and the rowi kiwi by first performing an error-correction step on the adapter-trimmed reads in tadpole from the BBtools toolsuite (Bushnell, 2014) and a kmer size of 31. We constructed the *de novo* assembly with the error-corrected reads using SOAPdenovo2 pregraph and contig (Luo et al., 2012), specifying a kmer size of 51, and otherwise using default parameters.

To construct the cross-species scaffolded assemblies we used the contig-level assemblies from above and scaffolded them either five times independently in the case of the beluga, or three times independently in the case of the rowi kiwi. For this, we constructed in-silico MP libraries using a modified version of the cross-species scaffolding pipeline (Grau, Hackl, Koepfli, & Hofreiter, 2018) and repeat-masked versions of the published non-beluga assemblies (narwhal, finless porpoise, bottlenose dolphin, sperm whale, minke whale) and published non-rowi assemblies (brown kiwi, spotted kiwi, emu). Repeats were masked based on the Genbank annotations.

In short, we constructed a fasta consensus sequence using a consensus base-call approach (-doFasta 2) in ANGSD v0.931 (Korneliussen, Albrechtsen, & Nielsen, 2014) and a minimum read depth of 3 (-minInddepth 3), minimum mapping quality of 25 (-minmapq 25), minimum base quality of 25 (-minq 25), and only considered reads that mapped uniquely to one location (-uniqueonly 1). We converted this fasta sequence into a pseudo-fastq sequence with a quality score of 40 for all covered bases using BBtools (Bushnell, 2014), as input for the seq-scripts pipeline.

From the consensus sequences, we constructed in-silico MP libraries with approximate insert sizes of 1kb, 2kb, 3kb, 5kb, 8kb, 10kb, 15kb, and 20kb using seq-scripts (https://github.com/thackl/seq-scripts), specifying a read length of 150bp and read depth of 100x. We scaffolded the contig-level beluga and rowi kiwi assemblies using SOAPdenovo2 map and scaff. To reduce the chances of mis-assembly by the longer in-silico MP libraries, we specified MP libraries of different insert sizes as different ranks in the SOAP config file. The shortest insert sizes had higher rankings; if a longer-insert library contradicted the shorter inserts, they were not used for scaffolding.

We assessed the final assembly quality in the form of contiguity (N50) and amount of missing data using QUAST v4.5 (Gurevich, Saveliev, Vyahhi, & Tesler, 2013).

### Sex-scaffold filtering

To exclude sex-linked scaffolds in downstream demographic and heterozygosity analyses, we determined the scaffolds that likely originated from the sex-chromosomes for each of the scaffolded assemblies (published assemblies and the cross-species scaffolded assemblies) used in this study. We found putative sex-chromosome scaffolds in the cetacean species by aligning all assemblies to the Cow X (Genbank accession: CM008168.2) and Human Y (Genbank accession: NC_000024.10) chromosomes, and putative sex-chromosome scaffolds in the palaeognath species by aligning all assemblies to the chicken (*Gallus gallus*) W (Genbank accession: CM000121.5) and Z (Genbank accession: CM000122.5) chromosomes. We did this using satsuma synteny v2.1 (Grabherr et al., 2010) and default parameters. We also removed scaffolds smaller than 10 kilobase pairs (kb) from all downstream analyses.

### Divergence estimates

To ensure comparability between the divergence estimates of our datasets, we calculated the autosome-wide divergence of our species of interest, either beluga or rowi kiwi, to the other species included in the study. To do this, we downloaded the raw reads for all the species (Supplementary table S3) and mapped them back to either the published beluga assembly or rowi kiwi assembly. We calculated the pairwise distance between species from the resultant mapped bam files and a consensus base call approach in ANGSD (-doIBS 2), and specifying the following parameters: -minq 25 -minmapq 25 -minind 4 -setMinDepthInd 5 -uniqueonly 1 -docounts 1 make a distance matrix (-makematrix 1), and only including autosomal scaffolds over 10kb in length (-rf).

### Demographic reconstruction

To determine the influence of (a) phylogenetic distance of the reference genome to the target species, (b) reference genome contiguity, and (c) the utility of cross-species scaffolded reference genomes on demographic reconstruction, we ran a Pairwise Sequentially Markovian Coalescent model (PSMC) (Li & Durbin, 2011) on each diploid genome, resulting in a total of twelve replicates for the beluga dataset and eight for the rowi kiwi dataset. We called diploid genome sequences using SAMtools and BCFtools v1.6 (Narasimhan et al., 2016), specifying a minimum quality score of 20 and minimum coverage of 10.

We ran PSMC specifying atomic intervals 4+25*2+4+6. Beluga PSMC outputs were plotted using a generation time of 32 years (Garde et al., 2015) and mutation rate of 1.65e-08 (Michael V. Westbury et al., 2019). To plot the rowi kiwi PSMC outputs, we calculated a mutation rate using the pairwise distance of the rowi kiwi to the brown kiwi (0.003123) and the formula pairwise distance × 2 / divergence time. We used a divergence time ∼3.8 Ma (De Cahsan & Westbury, 2020) which resulted in a mutation rate of 1.64×10^−9^ per year or 4.1×10^−8^ per generation assuming a generation time of 25 years (Weir, Haddrath, Robertson, Colbourne, & Baker, 2016).

### Genetic diversity

To determine the influence of (a) phylogenetic distance of the reference genome to the target species, (b) reference genome contiguity, and (c) the utility of cross-species scaffolded reference genomes on genetic diversity estimates, we estimated the autosome-wide heterozygosity of the beluga mapped to all twelve cetacean assemblies and the rowi kiwi mapped to all eight paleognath assemblies.

We calculated heterozygosity for each of the datasets using ANGSD. We estimated autosomal heterozygosity using allele frequencies (-doSaf 1), taking genotype likelihoods into account with the GATK algorithm (-GL 2), and specifying the following filters: only include sites with a read depth of at least 5 (-mininddepth 5), minimum mapping and base qualities of 30 (-minmapq 30, -minq 30), only include reads mapping uniquely to one location (-uniqueonly 1), only include reads where both read pairs map (-only_proper_pairs 1), only include autosomal scaffolds (-rf), and the extended adjust quality scores around indels parameter (-baq 2). Heterozygosity was computed from the output of this using realSFS from the ANGSD toolsuite, specifying 20 megabase pairs (Mb) windows of covered sites (-nSites).

We subsequently tested whether the results are consistent regardless of parameter and software selection using the beluga dataset mapped to the six published assemblies. We assessed the influence of parameter selection in ANGSD on heterozygosity estimates by computing heterozygosity using the procedure outlined above, but replacing the ‘extended adjust quality scores around indels’ parameter (-baq 2), with (i) adjust quality scores around insertion/deletions (indels) (-baq 1), (ii) no indel quality score adjustment (-baq 0), or (iii) extended adjust quality scores around indels (-baq 2), and adjust quality for reads with multiple mismatches to the reference (-C 50).

To assess whether software choice can impact results, we used two additional methods to compute heterozygosity of the beluga mapped to the six published assemblies: ROHan (Renaud, Hanghøj, Korneliussen, Willerslev, & Orlando, 2019), and the PSMC input diploid file (SAMtools/BCFtools).

In ROHan, we used default parameters to calculate autosome-wide levels of heterozygosity and runs of homozygosity (ROH). The default parameters specify a 1 MB window as being a ROH, if the window has an average heterozygosity of less than 1e-5. To calculate average autosome-wide heterozygosity from the diploid file used for the PSMC analysis, we used seqtk comp (https://github.com/lh3/seqtk).

### Inbreeding (runs of homozygosity)

As ROHan simultaneously outputs runs of homozygosity as well as autosome-wide levels of heterozygosity, we could evaluate how reference genome phylogenetic distance influences perceived inbreeding estimates using ROH. We did not retrieve any significant ROH in the beluga dataset using ROHan, so we were unable to investigate this further using the beluga data. We repeated the above ROHan analysis using the rowi kiwi mapped to all eight assemblies (both published and cross-species scaffolded).

## Results

### Mapping

Mapping results of the beluga raw reads to each cetacean assembly can be found in supplementary tables S4 and S5. Mapping results of the rowi kiwi raw reads to each paleognath assembly can be found in supplementary tables S6 and S7. As phylogenetic distance to the reference genome increases, there is a general trend of a decreasing number of unique reads mapping. This trend is not seen when mapping to the five and three cross-species scaffolded assemblies for beluga and rowi kiwi, respectively. However, less reads map to these assemblies than to the conspecific assemblies.

### Cross-species scaffolded *de novo* assemblies

Our contig-level beluga assembly had an N50 of ∼3.5 kb. The cross-species scaffolded assemblies were more contiguous, with N50s ranging from 283 kb to 614 kb (Supplementary table S8). However, these assemblies also had a lot of introduced missing data (16% - 18%). In comparison, the original assemblies for each species had N50s of 6.3 Mb - 122.2 Mb, and missing data rates of 0.5% - 6% (Supplementary table S1).

Our contig-level rowi kiwi assembly had an N50 ∼6.6 kb. The cross-species scaffolded assemblies were more contiguous, with N50s ranging from 1.9 Mb - 4.9 Mb (Supplementary table S9). These assemblies also had large amounts of introduced missing data (10% - 21%). However, this was comparable to the brown kiwi assembly with 14% data. Assembly contiguities were also more comparable to the published assemblies, which had N50s of 1.4 Mb - 5.7 Mb (Supplementary table S2).

### Demographic reconstruction

#### Beluga

With increasing phylogenetic distance of the reference genome, we see an incremental increase in deviation from the pattern obtained when mapping to the published beluga assembly (Fig 2A). However, we do not see an incremental change when using our five cross-species scaffolded assemblies as reference. Instead we see that all newly assembled genomes produce the same PSMC output. However, this output differs from the pattern obtained when mapping to the published beluga assembly (Fig 2B).

**Figure 2:**
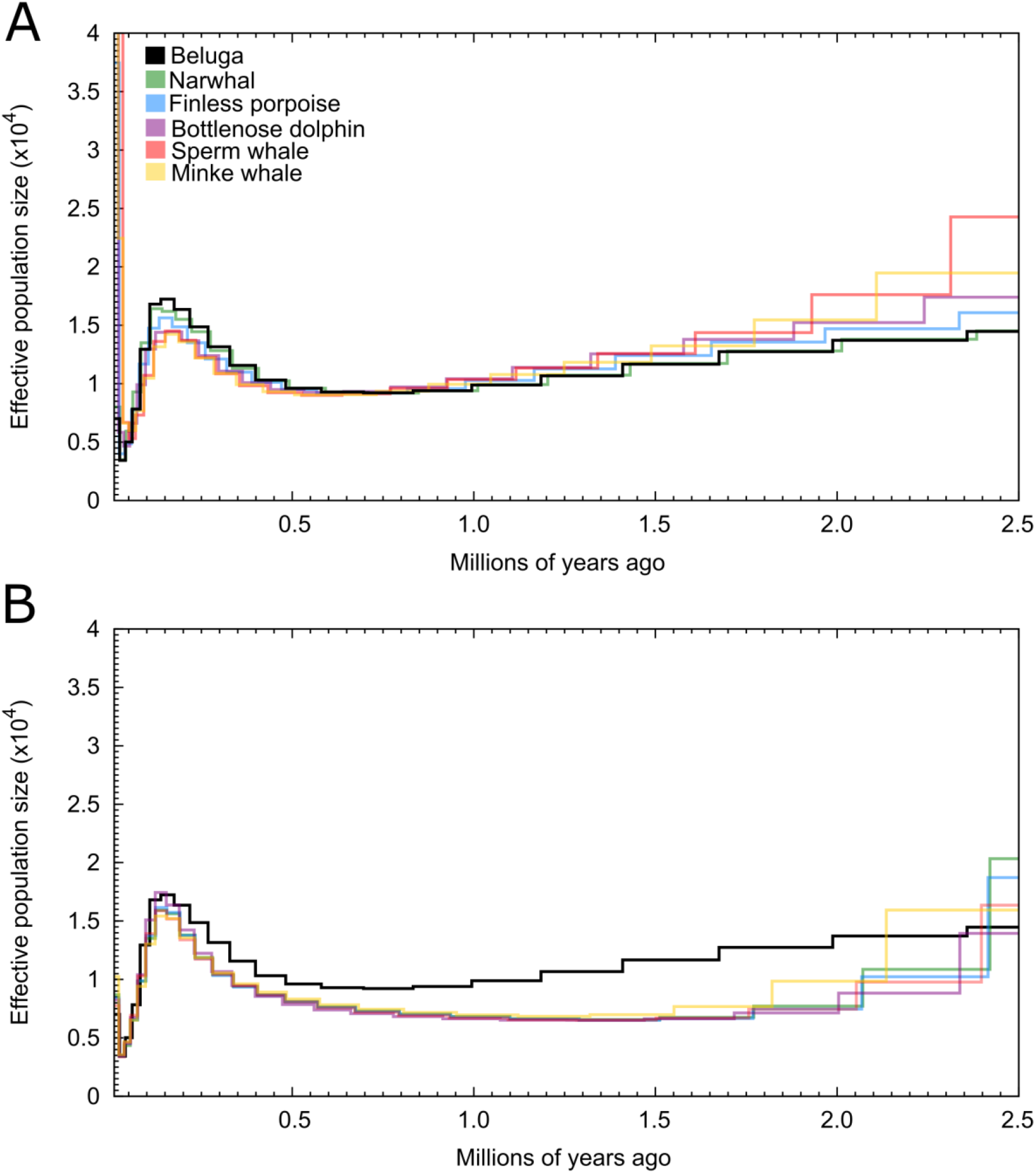
Beluga demographic history over the last 2.5 million years. Demographic trajectories in each panel represent genomes generated by mapping beluga reads to (A) assemblies of six different phylogenetically distant species (including a conspecific), colours indicate species of the reference genome, and (B) *de novo* assemblies constructed using cross-species scaffolding and the published beluga assembly, colours show the species used to scaffold the *de novo* beluga contig-level assembly.

When comparing PSMC results produced by mapping to the published beluga assembly, and by mapping to our *de novo* contig-level beluga assembly, we see a pattern of increase in N_e_ ∼500 thousand years ago (kya) followed by a decrease ∼150 kya. This is consistent between both assemblies. However, the values of effective population size (N_e_) are much lower when mapping to the *de novo* contig-level assembly (Supplementary fig S1).

#### Rowi kiwi

Unlike the beluga, the PSMC results of the rowi kiwi were vastly different when mapping to phylogenetic distant references compared to the published rowi assembly (Supplementary fig S2). However, we do see the incremental change as phylogenetic distance increases when mapping to the non-rowi assemblies. We investigated if there was a problem with the published rowi assembly by reassembling it using the published short-read and 3 kb mate-paired libraries (Sackton et al., 2019) with SOAPdenovo. Our reassembled rowi kiwi genome was much less contiguous than the published version (0.3 Mb vs 1.7 Mb) and had more missing data (12.7% vs. 1.6%). However, the PSMC produced when mapping to this assembly was much more consistent with what we would have expected based on the beluga results, and shows a demographic history similar to when mapping to the assemblies from the other three non-rowi kiwi species (Fig 3A). Hence, we only considered this re-assembly when assessing the inference of reference genome on demographic history results in the rowi kiwi. The results produced after mapping to the cross-species scaffolded rowi kiwi assemblies are much more similar to those from the re-assembled published rowi kiwi assembly (Fig 3B).

**Figure 3:**
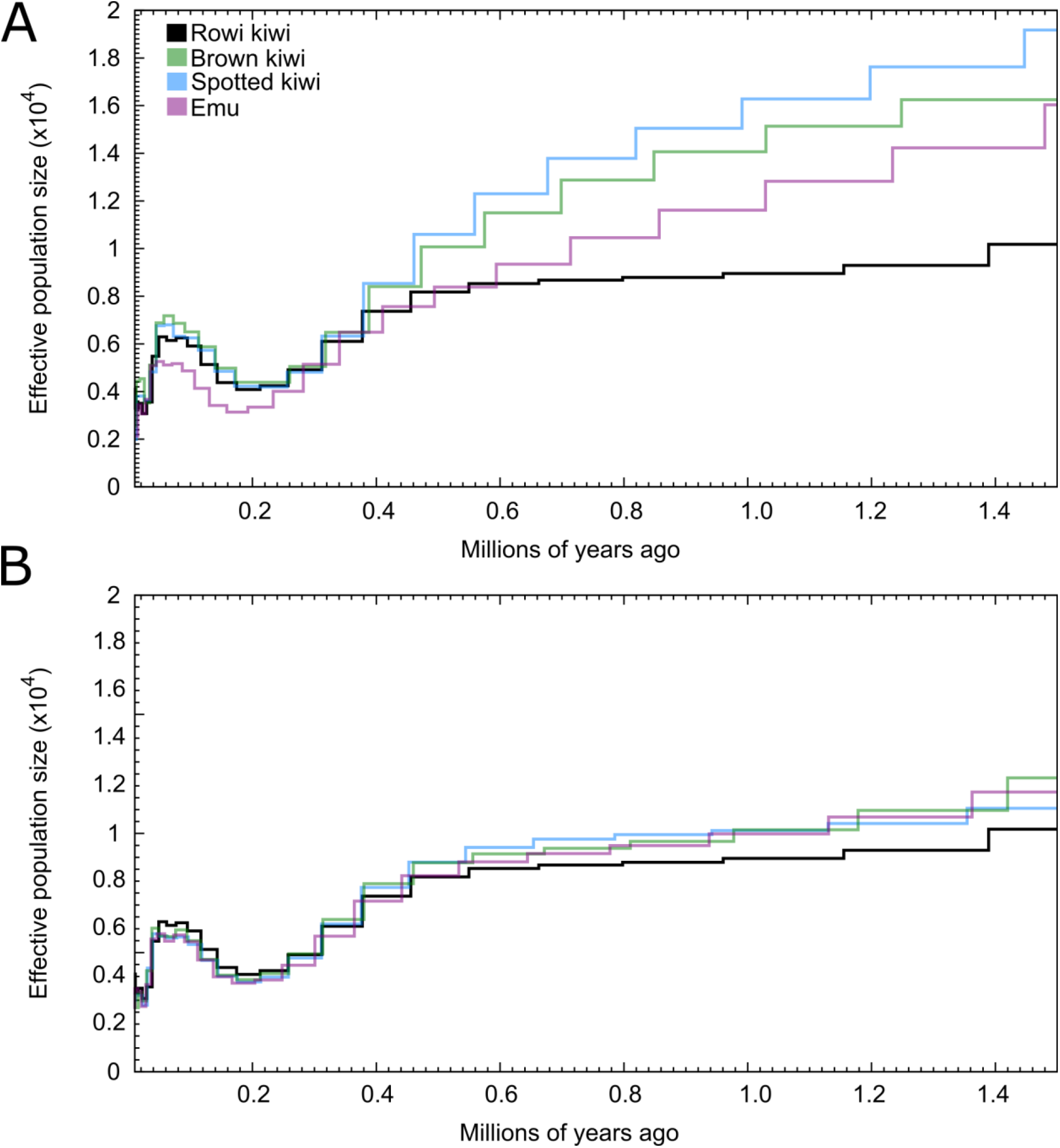
Rowi kiwi demographic history over the last 1.5 million years. Demographic trajectories in each panel represent genomes generated by mapping rowi kiwi reads to (A) assemblies of four different phylogenetically distant species (including our re-assembled rowi kiwi assembly), colours indicate species of the reference genome, and (B) *de novo* rowi kiwi assemblies constructed using cross-species scaffolding and our reassembled rowi kiwi assembly - colours show the species used to scaffold the *de novo* rowi kiwi contig-level assembly.

When comparing PSMC results produced by mapping to the re-assembled rowi genome, and by mapping to the contig-level assembly, we see similar general trajectories. However, the values of effective population size (Ne) are much lower when mapping to the *de novo* contig-level assembly as seen in the beluga (Supplementary fig S3).

### Genetic diversity

When using ANGSD, ROHan, and SAMtools/BCFtools, we see a general trend of increasing heterozygosity as reference genome phylogenetic distance increases. Which is also consistent when applying the alternative ANGSD parameter sets (i) -baq 1 instead of -baq 2, and (ii) -baq 0 instead of -baq 2 (Fig 4A,B, Supplementary figs S4 and S5, Supplementary tables S10 and S11). In contrast, when using ANGSD parameter set (iii) adjusted for reads with multiple mismatches to the reference (-C 50), we see a general trend of decreasing heterozygosity levels as phylogenetic distance increases (Supplementary fig S6).

**Figure 4:**
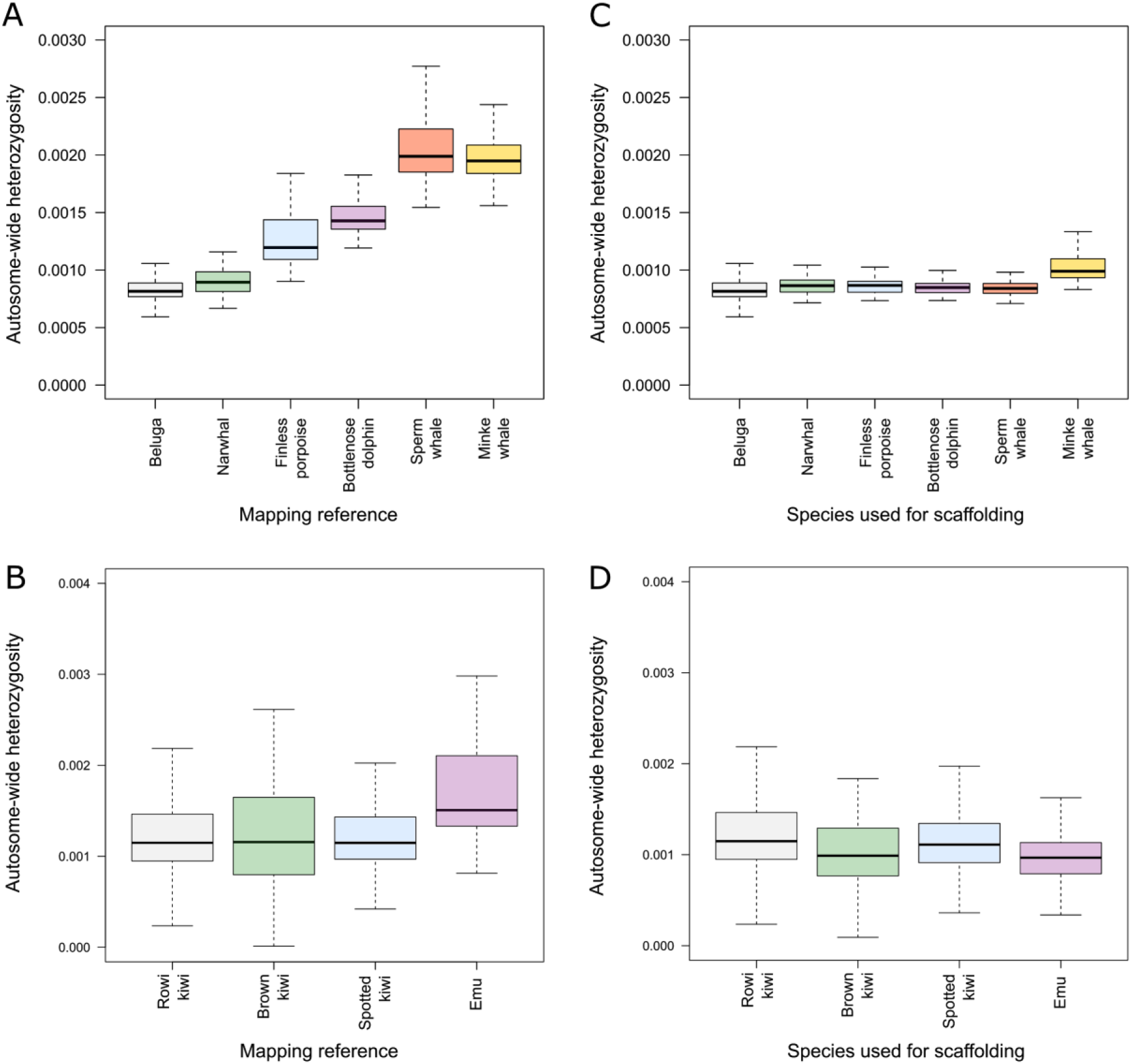
Autosome-wide heterozygosity estimates of the beluga and rowi kiwi mapped to different reference genomes. A single beluga individual was mapped to (A) six downloaded assemblies, and (B) a published beluga assembly and *de novo* beluga assemblies constructed using cross-species scaffolding. A single rowi kiwi individual was mapped to (C) our re-assembled rowi kiwi genome and the three downloaded non-rowi kiwi assemblies, and (D) *de novo* rowi kiwi assemblies constructed using either the published rowi kiwi mate-pair libraries or using cross-species scaffolding with mate-pair (MP) libraries constructed from each of the three non-rowi kiwi assemblies.

When using the cross-species scaffolded assemblies as reference genomes, we obtain results more comparable to those obtained when using the published conspecific assemblies as reference (Fig 4C,D). However, we do not see this when using SAMtools/BCFtools and the beluga dataset. Instead, we observe a decrease in heterozygosity relative to the published conspecific beluga assembly. The decrease is of a similar magnitude regardless of which cetacean species was used for scaffolding (Supplementary table S12).

The quality of the assembly also appears to play a role; higher genome-wide heterozygosity was estimated when mapping to the *de novo* contig-level beluga assembly compared to a scaffolded assembly (Supplementary fig S7). This same pattern was also seen when comparing the *de novo* contig-level rowi kiwi assembly to our reassembled version, but not compared to the published assembly (Supplementary fig S8).

### Inbreeding

When mapping the beluga reads to any reference genome (including the published conspecific beluga assembly), we did not uncover any ROH. When running ROHan on the rowi kiwi mapped to a published conspecific rowi kiwi assembly, we uncovered ROH, but not when mapping to any of the non-rowi kiwi assemblies (Table 2).

**Table 2:** Autosomal heterozygosity and runs of homozygosity (ROH) estimates of the rowi kiwi when mapped to a variety of different reference genomes. Reference genomes named ‘Rowi -’ are constructed using cross-species scaffolding and the species depicted after the hyphen.

| Reference genome | Global heterozygosity rate | Lower limit | Upper limit | Segments in ROH (%) | Avg. length of ROH (bp) |
| --- | --- | --- | --- | --- | --- |
| Rowi (published) | 0.00121 | 0.00111 | 0.00126 | 4.42 | 1,644,440 |
| Rowi (re-assembled) | 0.00105 | 0.00086 | 0.00123 | 1.57 | 1,076,920 |
| Brown kiwi | 0.00121 | 0.00107 | 0.00139 | 0.22 | 1,000,000 |
| Spotted kiwi | 0.00119 | 0.00108 | 0.00134 | 0.00 | 0 |
| Emu | 0.00177 | 0.00157 | 0.00193 | 0.00 | 0 |
| Rowi - brown kiwi | 0.00107 | 0.00097 | 0.00116 | 2.74 | 1,833,330 |
| Rowi - spotted kiwi | 0.00106 | 0.00098 | 0.00117 | 2.82 | 1,444,440 |
| Rowi - emu | 0.00099 | 0.00089 | 0.00107 | 3.28 | 1,558,140 |

## Discussion

Through a detailed comparison of results produced after mapping to multiple reference genomes from two unique datasets, we show that the choice of reference genome for mapping of short-read data of a target species can and does impact downstream evolutionary inferences. In general, as the phylogenetic distance of the reference genome increases, results become incrementally less reliable with regards to demographic history, genetic diversity, and inbreeding estimates.

With regards to demographic history analyses using PSMC, phylogenetic distance of the reference genome to the target species did not appear to affect the overall trajectories, but did result in relatively decreased N_e_ estimates. However, this only became apparent when using a reference genome more than 0.14% different to the target species (e.g. beluga vs finless porpoise) (Figs 2A, 3A). Based on these results, if a conspecific assembly is not available, using an assembly from a relatively closely-related species is unlikely to interfere with the overall demographic trajectory.

In contrast with the demographic results, the bias that reference-genome selection plays on genetic diversity estimates is more noticeable. Reference bias can cause heterozygous sites to be incorrectly called as homozygous for the reference allele (Brandt et al., 2015; Ros-Freixedes et al., 2018). However, we see a general increase in heterozygosity, as opposed to the expected decrease (Fig 4). Therefore, misalignments may be a larger factor in falsely calling heterozygous alleles as opposed to simply incorrect base calling. When applying a strict filter that corrects for reads with multiple mismatches to the reference genome (-C 50), it may be possible to eliminate increased heterozygosity due to misalignments (Supplementary figure S6). However, this is still associated with issues, as we observed a general decrease in heterozygosity due to putatively incorrect basecalls.

Phylogenetic distance driven reference bias was especially apparent when estimating ROH (Table 2). When mapping to a non-conspecific reference genome, we observed a complete loss of ROH, which would lead to the incorrect inference of no inbreeding in this individual. As we show that global heterozygosity rates increase as phylogenetic distance of the reference increases, this could artificially increase the heterozygosity level in ROH, making the ROH no longer observable.

One method we investigated for its putative ability to overcome these biases, without performing a traditional conspecific *de novo* assembly, is cross-species scaffolding. The biggest attraction of a cross-species scaffolded assembly over a traditional conspecific *de novo* assembly is that it only requires a single lane of Illumina sequencing, and an available assembly from a closely-related species. However, at least in the case of the beluga dataset, it can result in much more fragmented assemblies (Supplementary tables S8), and may therefore not always be applicable, especially when highly contiguous assemblies are required (e.g. for PSMC and ROH analyses). Nevertheless, using cross-species scaffolded assemblies as reference resulted in relatively reliable PSMC and ROH results for the rowi kiwi (Fig 3, Table 2), as well as reliable genetic results in all comparisons when using ANGSD (Fig 4). The reliability of these results may reflect that ANGSD uses genotype likelihoods to call heterozygosity (Korneliussen et al., 2014), as opposed to direct genotype calls, and therefore may be more reliable when heterozygous sites do not have the perfect near-50/50 allele ratios.

Despite the promising results when using cross-species scaffolded assemblies with our rowi kiwi dataset, PSMC results were less reliable using the beluga dataset. This could result from the low quality and highly-fragmented nature of the beluga cross-species scaffolded assemblies (Supplementary table S8). The inability to produce contiguous scaffolds like the rowi kiwi, may have arisen due to the highly fragmented nature of the beluga contig assembly, with an N50 of only ∼3.5 kb. To create a more contiguous final assembly, the N50 would need to be increased. Here, we implemented SOAPdenovo on a single library of random insert sizes. The use of different software (Butler et al., 2008) and lab protocols (Weisenfeld et al., 2014) to ensure the insert sizes are uniform may improve the contiguity of the final assembly, and make results reliant on highly contiguous data, such as PSMC, more reliable.

Although the assemblies are more fragmented, this does not mean they are completely devoid of information for the PSMC analysis. Results using the cross-species scaffolded assemblies still present the increase in N_e_ ∼500 kya and decrease ∼150 kya seen when using the published beluga assembly as reference, but with slightly decreased N_e_ values (Fig 2). Furthermore, when comparing PSMC results produced via mapping to the scaffold-level and contig-level assemblies, we also see a similar pattern of population size change, but with different values of N_e_ (Supplementary figs S1 and S3). This suggests that contiguity may not influence the pattern as much as the scale, and may still be useful for investigating relative changes in N_e_ rather than absolute values of N_e_ itself.

Our analyses uncovered a potential problem with the published rowi kiwi assembly. When comparing results mapped to the published assembly against non-conspecific assemblies, cross-species scaffolded assemblies, and a reassembly of the published data, we uncover large discrepancies in the results, especially in the PSMC results (Supplementary figs S2 and S3). As our assemblies all used the same published raw data, we suspect that these discrepancies resulted from miss-assemblies during the original *de novo* assembly process in Allpaths-LG (Butler et al., 2008). Although outside of the scope of the present study, these results show that caution should be exercised in reference genome selection for mapping assemblies; if multiple assemblies are available, it may be beneficial to test robustness of results against multiple reference genomes.

Taken together, our results show that demographic analyses of a single individual mapped to a phylogenetically distant reference genome may be considered reliable with regards to demographic trajectories (as in relative changes in N_e_, rather than absolute values of N_e_). However, the phylogenetic distance of the reference genome can lead to overestimation of heterozygosity and, in turn, underestimations of ROH. Finally, if no assembly from a suitably closely related species is available as a mapping reference, cross-species scaffolded assemblies appear to be a valid and likely more suitable option for evolutionary inference.

## Acknowledgements

The work was also supported by the Independent Research Fund Denmark | Natural Sciences, Forskningsprojekt 1, grant no. 8021-00218B to EDL.

## Author contributions

Conceptualization, MVW; Formal analysis, AP, MVW; Writing – Original Draft MVW; Writing – Review & Editing EDL, MVW; Funding Acquisition, EDL; Supervision, EDL, MVW.

