## Supplementary information for "Evaluating the role of reference-genome phylogenetic distance on evolutionary inference"

**Supplementary figures**

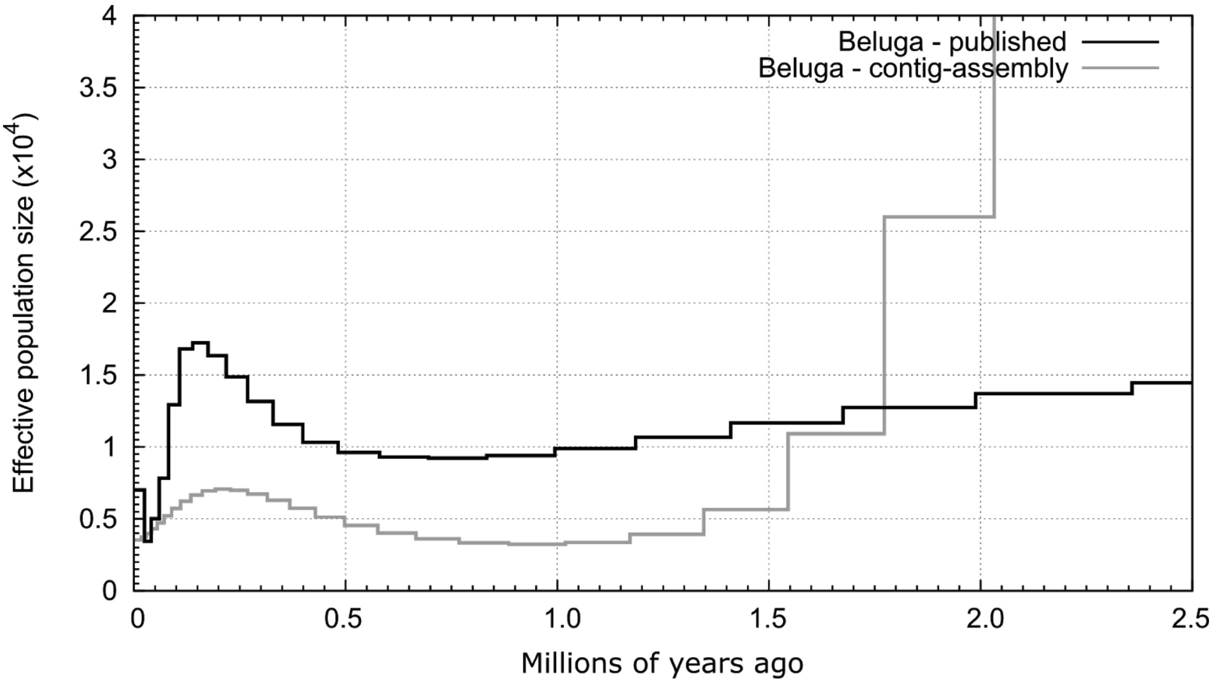


**Supplementary figure S1:** Demographic history over the last 2.5 million years of a single beluga individual mapped to a published scaffold-level beluga assembly and a *de-novo* contig-level beluga assembly.

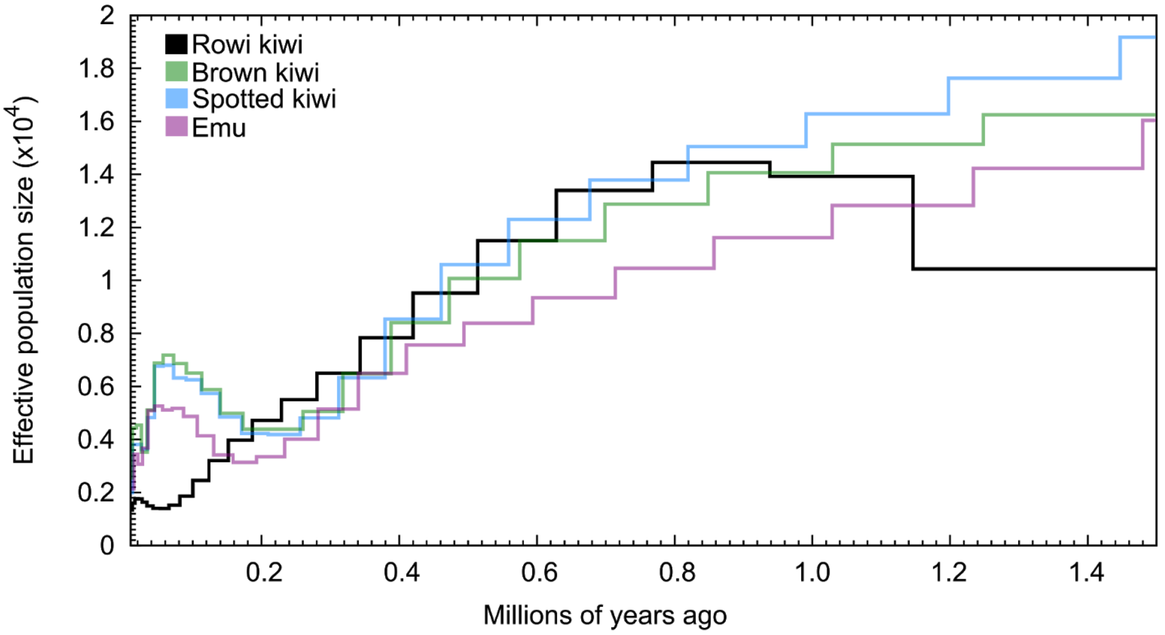


**Supplementary figure S2:** Rowi kiwi demographic history over the last 1.5 million years. Demographic trajectories represent those estimated based on genomes generated by mapping rowi kiwi reads to the published rowi kiwi assembly and three other paleognath assemblies - colours show the species used as mapping reference.

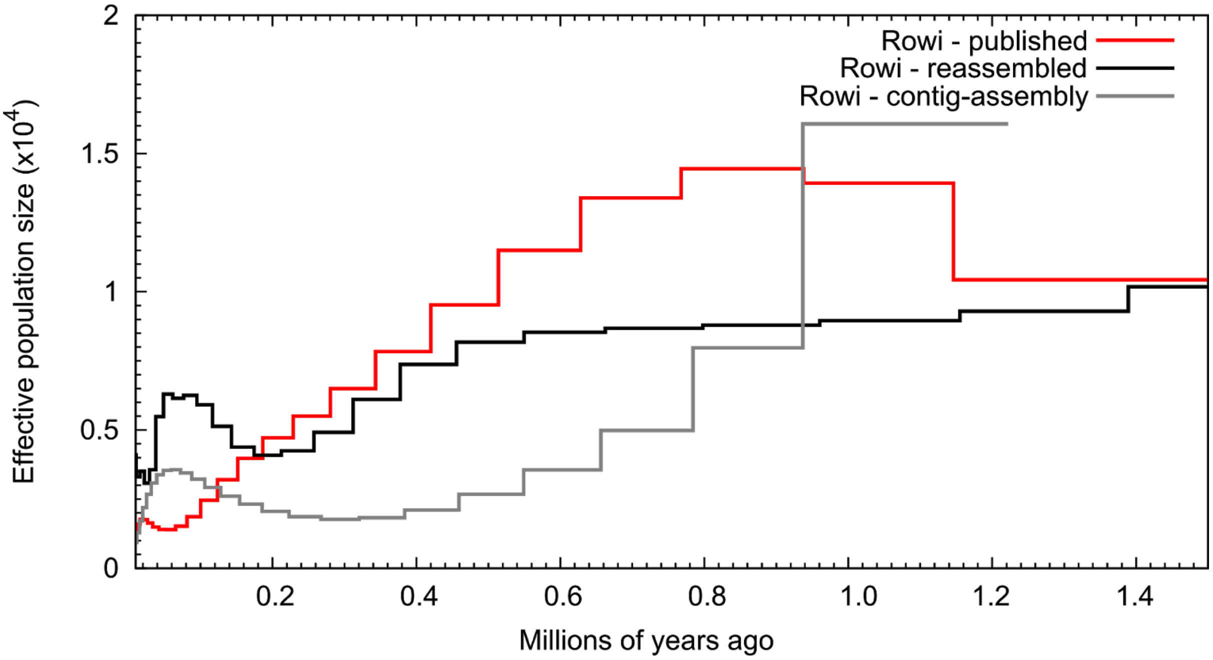


**Supplementary figure S3:** Rowi kiwi demographic history over the last 1.5 million years. Demographic trajectories represent those estimated based on genomes generated by mapping rowi kiwi reads to the published rowi kiwi assembly, our re-assembly of the published data, and our contig-level rowi kiwi assembly.

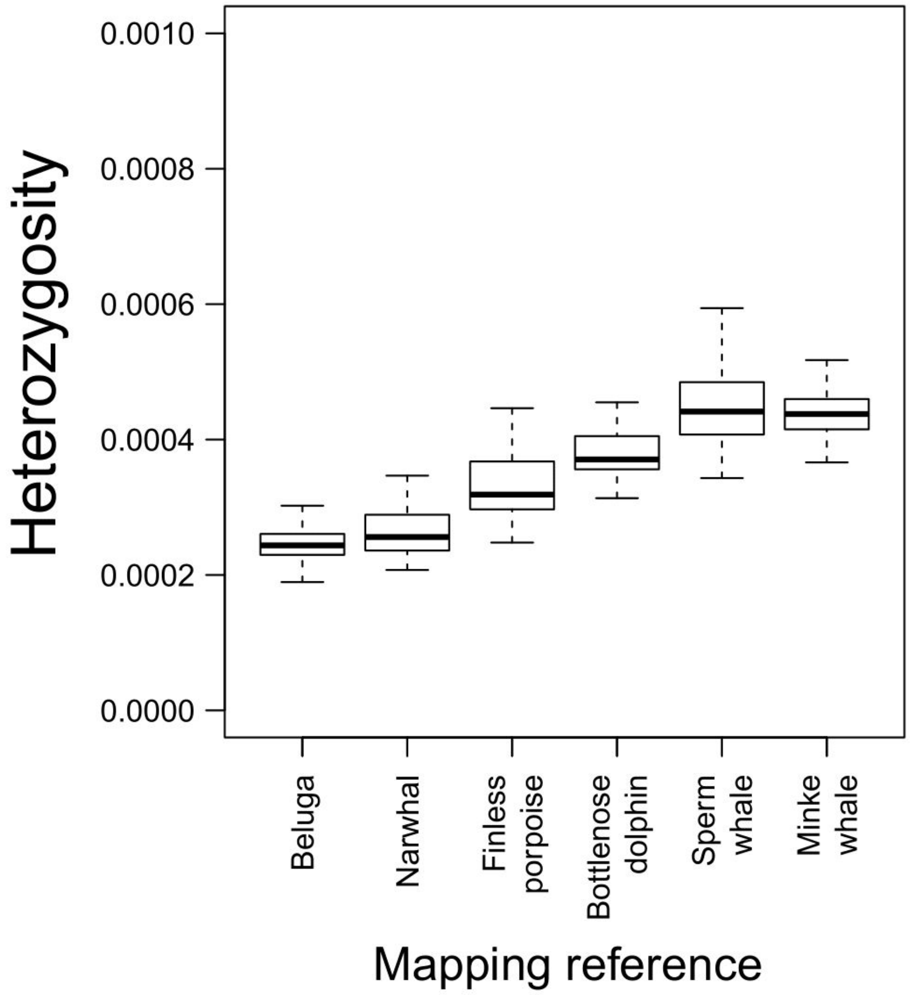


**Supplementary figure S4:** Autosome-wide heterozygosity results of a single beluga calculated in ANGSD using alternative parameter set i (baq 1) and various cetacean species as mapping reference.

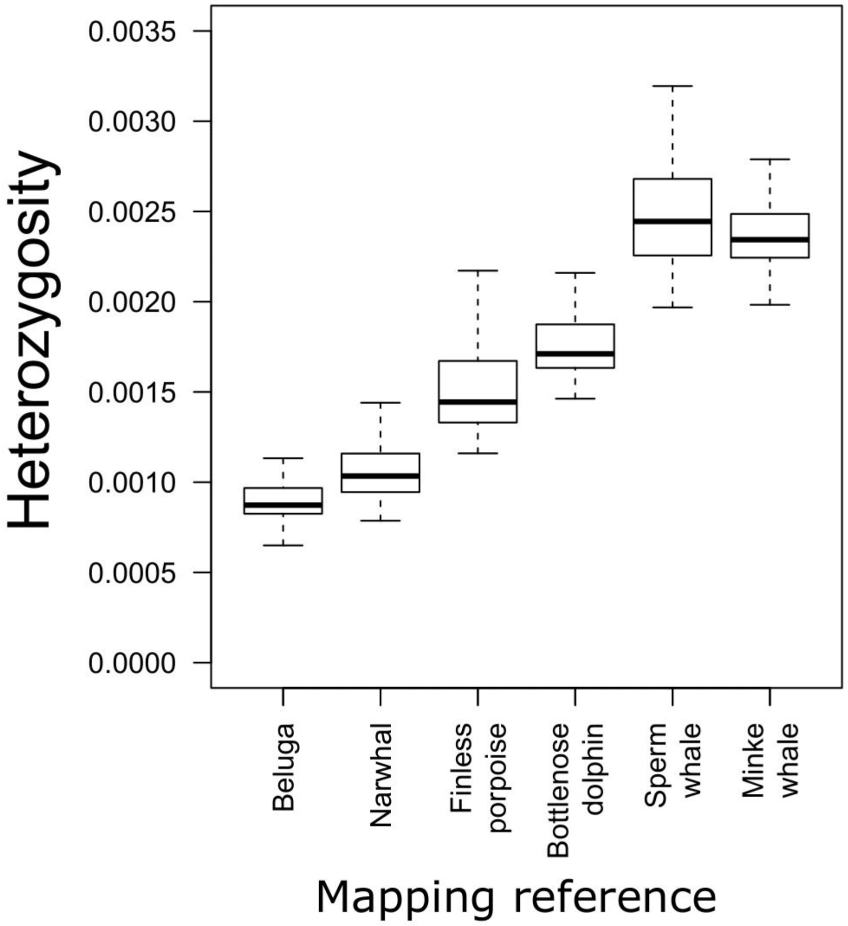


**Supplementary figure S5:** Autosome-wide heterozygosity results of a single beluga calculated in ANGSD using alternative parameter set ii (baq 0) and various cetacean species as mapping reference.

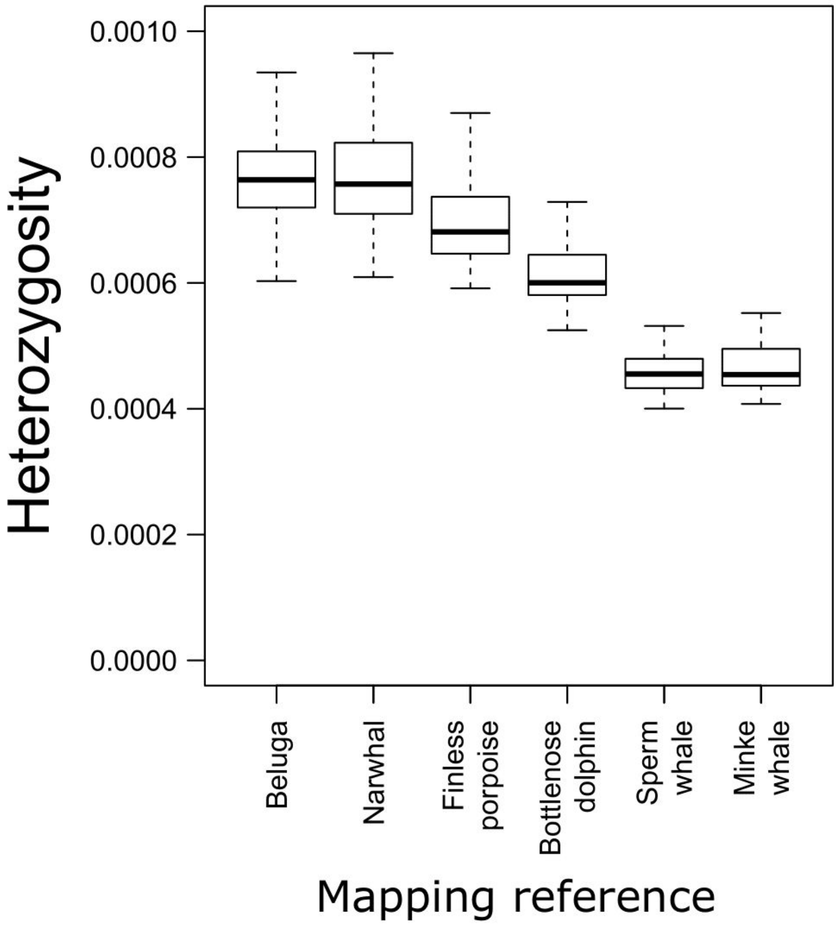


**Supplementary figure S6:** Autosome-wide heterozygosity results of a single beluga calculated in ANGSD using alternative parameter set iii (baq 2 and C 50) and various cetacean species as mapping reference.

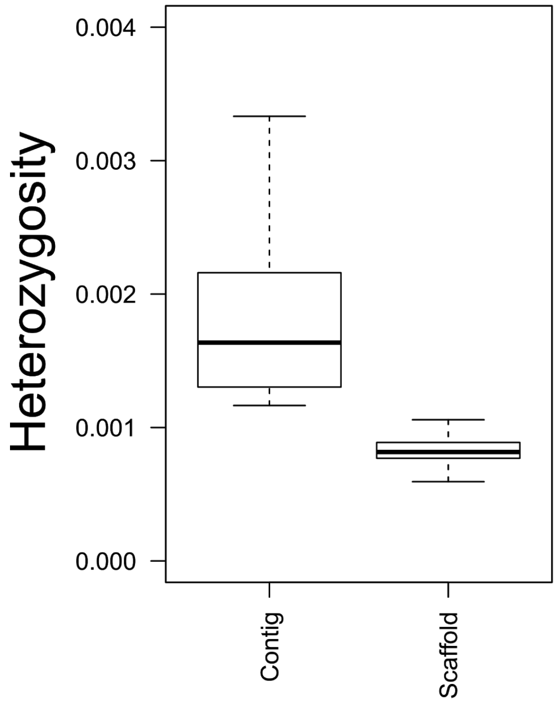


**Supplementary figure S7:** Autosome-wide heterozygosity results of a single beluga mapped to either our reconstructed contig-level beluga assembly or a downloaded scaffold-level beluga assembly.

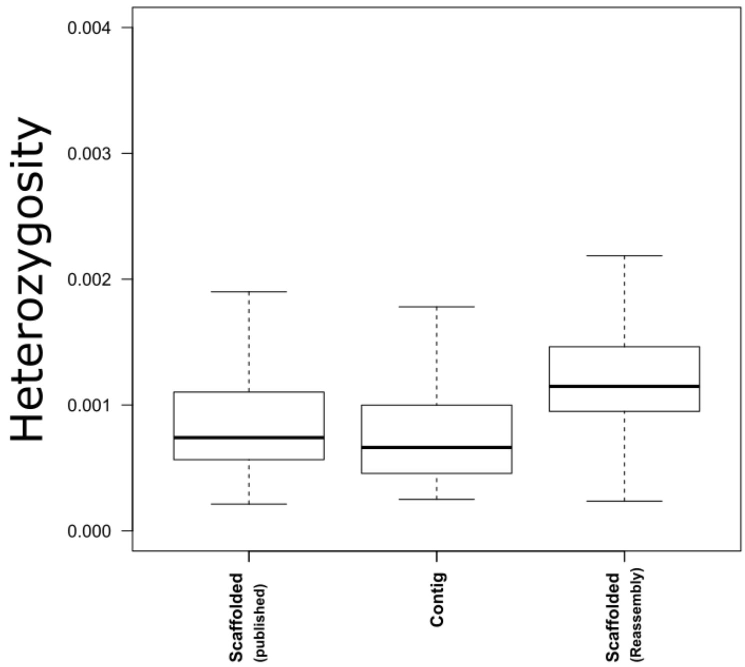


**Supplementary figure S8:** Autosome-wide heterozygosity results of a single rowi kiwi mapped to either a published scaffold-level rowi kiwi assembly, our reconstructed contig-level rowi assembly, or our scaffold-level rowi assembly which we re-assembled from published data.

**Supplementary tables**

**Supplementary table S1:** Assembly statistics for the various cetacean genomes downloaded from Genbank.

| Reference genome | Total length | N50 | Ns / 100 kb |
|---|---|---|---|
| Beluga | 2,362,774,659 | 31,183,418 | 1,517.93 |
| Narwhal | 2,355,574,979 | 107,566,389 | 486.89 |
| Finless porpoise | 2,284,628,084 | 6,341,296 | 715.51 |
| Bottlenose dolphin | 2,132,516,271 | 26,555,543 | 572.85 |
| Sperm whale | 2,512,149,402 | 122,182,240 | 4,117.25 |
| Minke whale | 2,423,101,967 | 13,026,147 | 5,968.16 |

**Supplementary table S2:** Assembly statistics for the various paleognath genomes downloaded from Genbank.

| Reference genome | Total length | N50 | N’s per 100 kb |
|---|---|---|---|
| Rowi | 1,228,902,913 | 1,672,276 | 1,570.8 |
| Brown kiwi | 1,523,972,539 | 5,679,020 | 13,974.76 |
| Spotted kiwi | 1,221,441,152 | 1,371,329 | 1,532.85 |
| Emu | 1192254075 | 3,305,683 | 1,106.95 |

**Supplementary table S3:** NCBI biosample accession codes for the raw reads used to calculate genome-wide divergences.

| Species | NCBI - Biosample |
|---|---|
| Narwhal | SAMN10519625 |
| Finless porpoise | SAMN02192673 |
| Bottlenose dolphin | SAMN09426418 |
| Sperm whale | SAMN07692275 |
| Minke whale | SAMD00016608 |
| Brown kiwi | SAMEA2554516 |
| Spotted kiwi | SAMN08476452 |
| Emu | SAMN08476457 |

**Supplementary table S4:** Mapping statistics of the beluga raw reads mapped to various cetacean assemblies downloaded from Genbank.

| Reference genome | Total reads | Read depth | Total mapped bp |
|---|---|---|---|
| Beluga | 531,650,735 | 34.49 | 79,239,197,768 |
| Narwhal | 529,266,471 | 34.17 | 78,711,565,548 |
| Finless porpoise | 503,630,565 | 33.44 | 74,418,529,882 |
| Bottlenose dolphin | 472,323,749 | 33.24 | 69,605,641,245 |
| Sperm whale | 445,863,847 | 30.67 | 64,799,289,636 |
| Minke whale | 446,967,504 | 31.57 | 65,029,200,042 |

**Supplementary table S5:** Mapping statistics of the beluga raw reads mapped to our newly constructed cross-species scaffolded beluga assemblies, each scaffolded using a different cetacean species.

| Reference genome | Cross-species MP libraries | Total reads | Read depth | Total mapped bp |
|---|---|---|---|---|
| Beluga-narwhal | Narwhal | 471,203,076 | 32.06 | 68,377,352,171 |
| Beluga-finless porpoise | Finless porpoise | 471,169,969 | 32.06 | 68,375,010,728 |
| Beluga-bottlenose dolphin | Bottlenose dolphin | 470,854,729 | 32.07 | 68,337,051,650 |
| Beluga-sperm whale | Sperm whale | 470,777,914 | 32.08 | 68,324,583,516 |
| Beluga-minke whale | Minke whale | 462,569,679 | 32.24 | 67,198,990,670 |

**Supplementary table S6:** Mapping statistics of the rowi kiwi raw reads mapped to various paleognath assemblies downloaded from Genbank.

| Reference genome | Total reads | Read depth | Total mapped bp |
|---|---|---|---|
| Rowi | 244,434,188 | 25.24 | 30,374,767,733 |
| Brown kiwi | 223,266,430 | 23.92 | 27,586,401,966 |
| Spotted kiwi | 242,085,076 | 25.08 | 30,006,509,667 |
| Emu | 195,821,276 | 20.87 | 22,979,468,373 |
| Rowi-reassembly | 237,542,646 | 24.71 | 29,247,194,839 |

**Supplementary table S7:** Mapping statistics of the rowi kiwi raw reads mapped to our newly constructed cross-species scaffolded rowi kiwi assemblies, each scaffolded using a different paleognath species.

| Reference genome | Cross-species MP libraries | Total reads | Read depth | Total mapped bp |
|---|---|---|---|---|
| Rowi-brown kiwi | Mantelli | 233,627,179 | 24.60 | 28,700,071,645 |
| Rowi-spotted kiwi | Haasti | 234,757,229 | 24.60 | 28,840,738,462 |
| Rowi-Emu | Emu | 232,361,399 | 24.68 | 28,574,913,977 |

**Supplementary table S8:** Assembly statistics for our newly constructed beluga assemblies, each scaffolded using cross species scaffolding with a different cetacean species.

| Reference genome | Cross-species MP libraries | Total length | N50 | Ns / 100 kb |
|---|---|---|---|---|
| Beluga-narwhal | Narwhal | 2,536,530,167 | 613,685 | 17,030.78 |
| Beluga-finless porpoise | Finless porpoise | 2,471,430,825 | 348,434 | 15,895.64 |
| Beluga-bottlenose dolphin | Bottlenose dolphin | 2,502,821,192 | 283,481 | 16,411.77 |
| Beluga-sperm whale | Sperm whale | 2,509,669,412 | 317,856 | 18,112.01 |
| Beluga-minke whale | Minke whale | 2,533,169,536 | 378,296 | 17,255.77 |

**Supplementary table S9:** Assembly statistics for our newly constructed rowi kiwi assemblies, each scaffolded using cross-species scaffolding with a different paleognath species.

| Reference genome | Cross-species MP libraries | Total length | N50 | N's per 100 kb |
|---|---|---|---|---|
| Rowi-brown kiwi | Brown kiwi | 1,574,166,636 | 4,887,347 | 21,111.41 |
| Rowi-Spotted kiwi | Spotted kiwi | 1,400,063,207 | 1,935,686 | 10,727.33 |
| Rowi-Emu | Emu | 1,385,314,252 | 1,887,596 | 9,620.41 |

**Supplementary table S10:** Heterozygosity and runs of homozygosity estimates of the beluga mapped to assemblies from various cetacean species. Calculated using ROHan.

| Reference genome | Beluga | Narwhal | Finless porpoise | Bottlenose dolphin | Sperm whale | Minke whale |
|---|---|---|---|---|---|---|
| Global het rate: | 0.000871 | 0.000952 | 0.001101 | 0.001272 | 0.001783 | 0.001711 |
| Lower limit | 0.000788 | 0.000869 | 0.001010 | 0.001173 | 0.001635 | 0.001573 |
| Upper limit | 0.000959 | 0.001035 | 0.001195 | 0.001368 | 0.001940 | 0.001851 |
| Segments in ROH(%) | 0 | 0 | 0 | 0 | 0 | 0 |
| Avg. length of ROH | 0 | 0 | 0 | 0 | 0 | 0 |

**Supplementary table S11:** Heterozygosity estimates of the beluga mapped to assemblies from various cetacean species. Calculated from the PSMC diploid input file.

| Reference genome | Total no of sites | Heterozygous sites | Heterozygosity rate |
|---|---|---|---|
| Beluga | 2,270,644,945 | 1,822,242 | 0.000803 |
| Narwhal | 1,689,332,137 | 1,436,141 | 0.000850 |
| Finless porpoise | 1,932,136,360 | 1,920,441 | 0.000994 |
| Dolphin | 1,642,645,105 | 1,874,404 | 0.001141 |
| Sperm whale | 1,432,735,888 | 1,961,116 | 0.001369 |
| Minke whale | 1,542,481,560 | 2,088,348 | 0.001354 |

**Supplementary table S12:** Heterozygosity estimates of the beluga mapped to our newly constructed beluga genomes, each scaffolded using a different cetacean species. Calculated from the PSMC diploid input file.

| Reference genome | Cross-species MP libraries | Total no of sites | Heterozygous sites | Heterozygosity rate |
|---|---|---|---|---|
| Beluga | NA | 2,270,644,945 | 1,822,242 | 0.000803 |
| Beluga-narwhal | Narwhal | 1,985,070,058 | 1,485,537 | 0.000748 |
| Beluga-finless porpoise | Finless porpoise | 1,983,218,243 | 1,475,312 | 0.000744 |
| Beluga-bottlenose dolphin | Bottlenose dolphin | 1,974,005,041 | 1,454,709 | 0.000737 |
| Beluga-sperm whale | Sperm whale | 1,964,210,075 | 1,432,092 | 0.000729 |
| Beluga-minke whale | Minke whale | 1,978,738,668 | 1,673,527 | 0.000846 |
